# Reflected Stochastic Differential Equation Models for Constrained Animal Movement

**DOI:** 10.1101/152017

**Authors:** Ephraim M. Hanks, Devin S. Johnson, Mevin B. Hooten

## Abstract

Movement for many animal species is constrained in space by barriers such as rivers, shore-lines, or impassable cliffs. We develop an approach for modeling animal movement constrained in space by considering a class of constrained stochastic processes, reflected stochastic differential equations. Our approach generalizes existing methods for modeling unconstrained animal movement. We present methods for simulation and inference based on augmenting the constrained movement path with a latent unconstrained path and illustrate this augmentation with a simulation example and an analysis of telemetry data from a Steller sea lion *(Eumatopias jubatus)* in southeast Alaska.

## 1 Introduction

The movement of animals and humans is a fundamental process that drives gene flow, infectious disease spread, and the flow of information and resources through a population (Hanks and Hooten, 2013; Coulon et al., 2006; Hooten et al., 2007; Scharf et al., 2015). Movement behavior is complex, often exhibiting directional persistence, response to local environmental conditions, dependence between conspecifics, and changing behavior in time and space. While technological advances have allowed movement (telemetry) data to be collected at high resolution in time and space, most movement data still exhibit non-negligible observation error, requiring latent variable approaches, such as hidden Markov models (HMMs) or Bayesian hierarchical models (BHMs) to provide inference for movement parameters. The field of movement ecology is broad and growing; many different modeling approaches have been proposed for different species exhibiting different behaviors (e.g., Hooten et al., 2017). A majority of attempts to model movement stochastically rely on unconstrained stochastic processes, with positive probability of movement to any region in space (typically 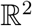). This continuous-space assumption is realistic for many species, but is clearly violated for others, such as marine animals swimming near shorelines (Bjørge et al., 2002; Johnson, London, Lea and Durban, 2008; Small et al., 2005) or ants constrained to walk inside the confines of a nest (Mersch et al., 2013; Quevillon et al., 2015). In addition, measurement error on telemetry data often results in biologically impossible recorded animal locations, such as a seal being located miles inland, or two successive ant locations being separated by an impassible wall.

We consider spatially-constrained animal movement, where an animal can only be present within a known subset 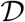 of 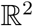. To illustrate the need for movement models constrained by space, we consider movement of a Stellar sea lion *(Eumatopias jubatus)*, a marine mammal that stays entirely in the water or hauled-out on the shoreline. Figure 1 shows telemetry data obtained using the ARGOS system (ARGOS, 2015) from one sea lion over a thirty-day observation period from December 6, 2010 to January 5, 2011. Stellar sea lions have experienced recent fluctuations in population size, and could be threatened by disease, increased fishing in Northern waters, and other factors (Dalton, 2005). Understanding where sea lions spend time can inform species management decisions and fishing regulations off the coast of Alaska. Telemetry data provide a natural approach to studying Stellar sea lion space use.

**Figure 1:**
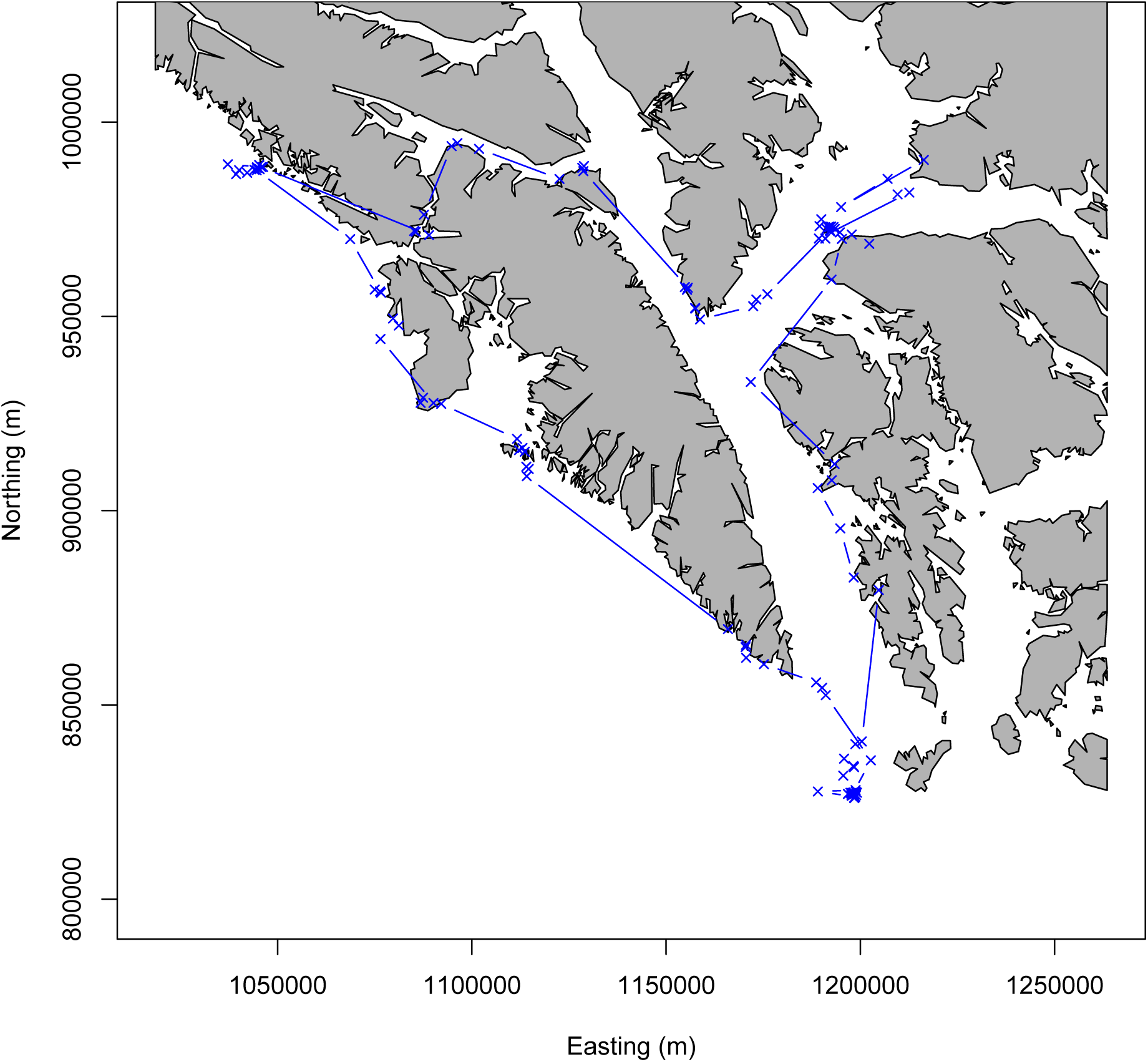
Sea Lion Telemetry Data. Telemetry data from 30 days of observation of a sea lion (*Eumatopias jubatus*) in southeast Alaska, obtained using the ARGOS system.

Remote tracking of marine mammals is challenging, because common tracking systems (such as GPS) are impeded by water. While the sea lion is always either in the water or hauled out meters from the water’s edge, many of the telemetry locations are kilometers inland (Figure 1). If a movement model were fit to the data without accounting for the constraint that the sea lion remain within water at all times, the posterior distribution of paths the animal could have taken would overlap land. This may lead to biased inference for space use or resource selection of pinnipeds (Brost et al., 2015), which could, in turn, lead to inefficient species management decisions. Additionally, inference without considering the spatial constraint (for example, the need to go around an island between telemetry observations) could lead to biased estimates for parameters governing animal movement.

Statistical inference for constrained movement is computationally challenging, because the spatial constraint 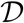 often makes the evaluation of density functions only possible numerically. We present an approach for modeling constrained animal movement based on reflected stochastic differential equations (RSDEs), which have been used to model constrained processes in many fields. To implement our approach, we present a Markov chain Monte Carlo (MCMC) algorithm for sampling from the posterior distribution of model parameters by augmenting the constrained process with an unconstrained process. We illustrate our approach through a simulation example, and an application to telemetry data from the sea lion shown in Figure 1.

## 2 Modeling Constrained Movement With Reflected Stochastic Differential Equations

Stochastic differential equation (SDE) models are popular stochastic process models for animal movement (Brillinger et al., 2002; Brillinger, 2003; Johnson, Thomas, Ver Hoef and Christ, 2008; Preisler et al., 2013; Russell et al., 2017). Brillinger (2003) considered simulation of animal movement under a constrained RSDE model, but did not consider inference under such a model. We develop a class of SDE models that can capture a wide range of movement behavior, and then propose approaches for simulation and inference under this class of models.

### 2.1 Modeling Observational Error

In general, we assume that we observe animal locations s*_t_*, *t* ∊ {*τ*_1_, *τ*_2_, …, *τ_T_*} at *T* distinct points in time {*τ_t_*, *t* = 1, …, *T*}. We assume that the locations are in 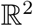, with 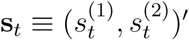 representing the observed location at time *τ_t_*. The extension to higher dimensions (e.g., three-dimensional space) is straightforward. The observations are assumed to be noisy versions of the true animal location 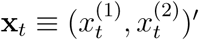 at time *τ_t_*, with observation error distribution

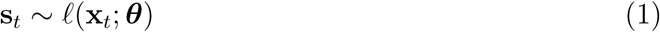

where *θ* contains parameters controlling the distribution of observations centered at the true location. We begin by leaving this observation error distribution unspecified, and develop a general framework for inference, then apply a specific class of models to the sea lion telemetry data.

To allow for switching between notation for discrete and continuous time processes, we adopt the following convention for subscripts. A greek letter in the subscript implies an observation in continuous time, with x*_τ_* being the location of the animal at time *τ*. We use a standard Latin letter in the subscript to index a set of locations at a discrete set of times; thus 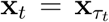 represents the individual’s location at time *τ_t_*. We also adopt the notation that x_s:t_ ≡ {x*_s_*, x*_s_*_+1_, …, x_*t*−1_, x*_t_*} represents the set of (*t* − *s* + 1) observations in discrete time between the sth and tth observations (inclusive) in the sequence.

In the next Section, we will develop a model for movement based on an approximate solution to an SDE. As this approximation operates in discrete time, with a temporal step size of *h*, the approximation yields latent animal locations x*_τ_* : *τ* ∊ {0, *h*, 2*h*, …, *Th*} at discrete times.

### 2.2 A general SDE model for animal movement

We first consider the unconstrained case 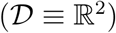, and then consider constrained processes. A class of SDE models that can capture a wide range of movement behavior are expressed as follows. Let the individual’s position at time *τ* be x*_τ_* and define v_*τ*_ to be the individual’s true velocity at time *τ*

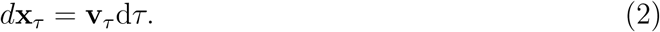

This differential equation may be equivalently written as an integral equation (e.g., Hooten and Johnson, 2017)

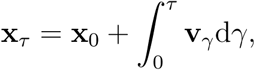

however, we adopt the differential equation form throughout this section.

By modeling the time derivative of an individual’s velocity, we focus on modeling acceleration, or, equivalently, the force applied to an individual animal over time. This provides a natural framework for modeling intrinsic and extrinsic forces applied to a moving animal. Consider the following SDE model for the time derivative of velocity

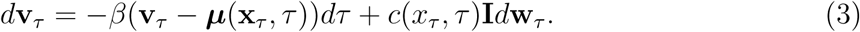

In (3), *β* is an autocorrelation parameter, *μ*(x*_τ_*, *τ*) is a function specifying the vector-valued mean direction of movement (drift), perhaps as a function of time *τ* or current location x*_τ_*, 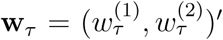 is a vector of two independent standard Brownian motion processes, **I** is the 2 × 2 identity matrix, and *c*(x*_τ_*, *τ*) is a scalar function controlling the magnitude of the stochastic component of (3).

Several existing models for animal movement fit into the general framework defined by (2)-(3). For example, the continuous-time correlated random walk model developed by Johnson, London, Lea and Durban (2008), with a constant drift *μ*, is obtained by setting *μ*(x*_τ_*, *τ*) = *μ* and assuming constant stochastic variance across time *c*(x*_τ_*, *τ*) = *σ*. Johnson, London, Lea and Durban (2008) also consider a time-varying drift parameter by modeling *d*v*_τ_* as the sum of two stochastic processes similar to those in (3), operating on different time scales.

As a second example, the potential function approach to modeling animal movement (Brillinger et al., 2001, 2002; Preisler et al., 2004, 2013) results from specifying the drift function as *μ*(x*_τ_*, *t*) = − ∇*H*(x*_τ_*), the negative gradient of a potential surface *H*(*x*), which is a scalar function defined in 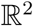. In the overdamped case where *β* → ∞, and when the stochastic variance is constant over time and space (*c*(x*_τ_, τ*) = *σ*), the SDE (3) reduces to

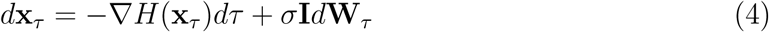

See Brillinger et al. (2001) for details. The velocity-based movement model of Hanks et al. (2011) results from taking a discrete (Euler) approximation to the SDE in (4).

As a third example, the spatially-varying SDE approach of Russell et al. (2017) for modeling spatial variation in motility (overall rate of speed) and directional bias could be approximated by setting *c*(x*_τ_*, *τ*) = *σm*(x*_τ_*) and 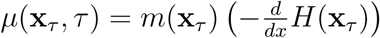, where *H*(x) is a potential function as in Brillinger (2001), and *m*(x*_τ_*) is a spatially-varying motility surface that acts by dilating or compressing time, as is done by Hooten and Johnson (2017) using a time warping function. While Russell et al. (2017) allow this motility or time-dilation surface to vary across space, Hooten and Johnson (2017) allowed their warping function to vary across time to capture time-varying movement behavior in which individuals exhibit periods of little or no movement interspersed with periods of higher activity.

### 2.3 Numerically approximating constrained SDEs

We consider simulation of a constrained SDE and describe a related approach for inference. The model in (2)-(3) is a semi-linear Ito SDE (e.g., Allen, 2007), and in some cases, such as the CTCRW of Johnson, London, Lea and Durban (2008) and the potential function approach of Brillinger et al. (2001), closed form solutions are available for this transient distribution without spatial constraints. However, when movement is constrained to occur within a fixed spatial domain 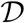, no closed form for the general transient distribution exists. Thus, we consider numerical approximations to the solution of the SDE, both without the spatial constraint and using modified approximations that account for the spatial constraint 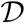.

The simplest and most common numerical approximation to the solution to the SDE (2)-(3) is the Euler-Maruyama scheme, which results from a first-order Taylor series approximation (e.g., Kloeden and Platen, 1992). Given a temporal step-size of *h*, the Euler-Maruyama iterations are

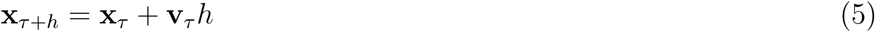

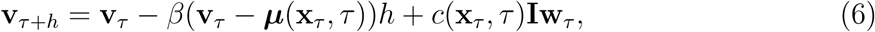

where 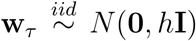. The numerical approximation in (5)-(6) is known to be of strong order 1/2 (Kloeden and Platen, 1992). Russell et al. (2017) use this Euler-Maruyama numerical procedure to specify an approximate statistical model for spatially-varying movement behavior of ants.

#### 2.3.1 A Two-Step Higher Order Procedure

Brillinger (2003) notes that, for constrained SDEs, the Euler-Maruyama scheme may require a very fine temporal discretization to result in realistic paths, and recommends higher-order numerical schemes be used. One modification of the above Euler-Maruyama procedure involves replacing the velocity v*_τ_* with a first difference approximation. This is similar to the approach taken in Runge-Kutta procedures for solving partial differential equations (e.g., Cangelosi and Hooten, 2009; Wikle and Hooten, 2010; Cressie and Wikle, 2011). From (5), note that v*_τ_* = (x*_τ_*_+*h*_ − x*_τ_*)/*h*. Substituting this expression for v*_τ_* into (6) gives

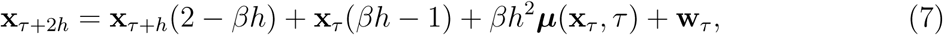

Where 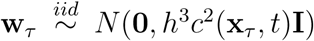 This numerical procedure has three main benefits relative to the Euler-Maruyama approach. First, the resulting solution to the unconstrained SDE is an approximation of strong order 1 (Kloeden and Platen, 1992) and thus provides a more accurate approximation to the continuous-time solution than does the Euler-Maruyama procedure. Second, this procedure removes the latent velocity v*_τ_* from the probability distribution, which simplifies the transition densities to only rely on animal locations at the two previous time points. Russell et al. (2017) used the Euler-Maruyama approach to motivate a statistical model, and treated v*_τ_* as latent variables to be estimated. The two-step procedure in (7) removes the need to make inference on the latent v. Third, removing the latent velocity from the approximation simplifies the solution in the presence of a spatial constraint 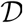, because the velocity is not constrained, but the animal’s position is constrained to occur within 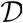.

#### 2.3.2 Reflected Stochastic Differential Equations for Animal Movement

The SDE in (2)-(3), whose solution is approximated by (7), is not constrained to occur within 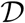. One theoretical approach to constructing a constrained process is to consider a process k*_τ_* that is defined as the minimal process required to keep x*_τ_* within 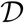. Thus, we modify (2)-(3) to obtain the constrained stochastic process

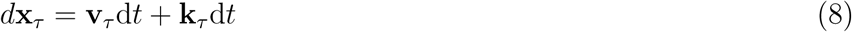

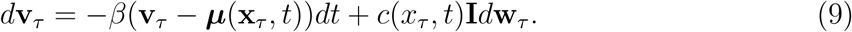

This approach is a so-called “reflected” stochastic differential equation (RSDE, e.g., Lépingle, 1995; Grebenkov, 2007; Dangerfield et al., 2012), a generalization of reflected Brownian motion. In reflected Brownian motion, a Brownian trajectory is reflected when it encounters the boundary 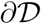 of the domain 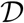. While there are many theoretical results for reflected Brownian motion, we note that the SDE in (2)-(3) is a variation on integrated Brownian motion, and therefore results for reflected Brownian motion are not directly applicable here.

The process k*_τ_* is defined as the minimal process required to restrict x*_τ_* to be within 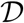, and can be described by considering a unit vector n(x) that points toward the interior of 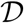 orthogonal to 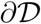 at x. Then this minimal process is defined as

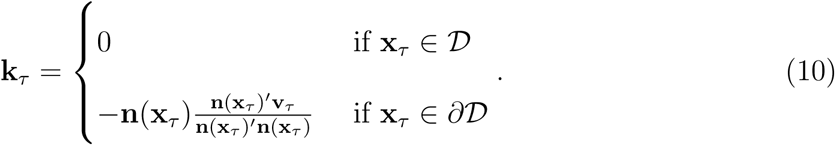

Under this specification, when an individual encounters the boundary 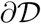, the process k*_τ_* nullifies the component of the individual’s velocity that would carry it out of 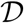, and the individual’s velocity becomes parallel to the boundary 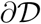 until acted upon by other forces (such as w*_τ_*). k*_τ_* is defined when 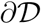 admits an orthogonal vector **n**, which is true for smooth boundaries 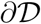. A natural way to define 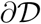 is as a polygon, which is piece-wise continuous. In this setting, n would be undefined at polygon vertices, but for a fine temporal resolution, the latent process x will rarely or never directly encounter the vertices.

The numerical solution (approximation) to such a constrained SDE (8)-(9) can be obtained in one of two ways. The most common approach (e.g., Lépingle, 1995; Grebenkov, 2007; Dangerfield et al., 2012) is to consider a *projected* version of a numerical solution to the unconstrained SDE. This corresponds to the projection approach proposed by Brillinger (2003) for a simpler SDE, who also proposes two other schemes for constraining x*_τ_* to remain within 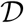. We do not consider these other schemes here, but make note of them in the Discussion.

In a projected approach to solving the RSDE, the two-time-step numerical procedure in (7) is modified by augmenting the solution x*_τ_* to the constrained SDE with an unconstrained process **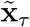** that may occur outside 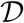, as follows. Conditioned on the constrained process at previous times x_1_:(*τ*+*h*), the distribution of the unconstrained process 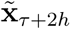is given by (7), with

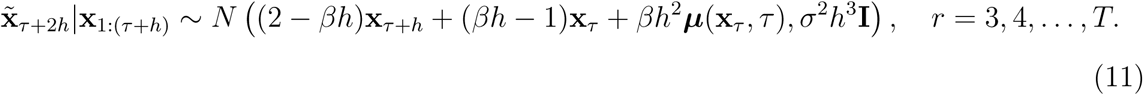

Any simulated animal location 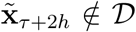 that falls outside of the spatial region 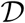 is projected onto the nearest location 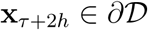 on the spatial boundary

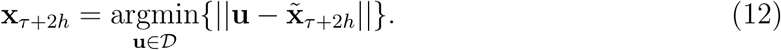

This results in a computationally efficient approach to simulating sample paths from the constrained SDE in (8)-(9), as the boundary 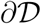 can be approximated as a polygon, and fast algorithms can be specified for projection of a point outside of 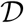 onto the polygonal boundary 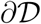. Pseudo-code for simulation of the RSDE in (8)-(9) for a given temporal step-size *h* is given in Appendix A, and R code to implement this approach is available upon request.

## 3 Inference on RSDE Model Parameters

We now consider inference on the movement parameters *θ* ≡ (*β*, *σ*^2^)′ from observed telemetry data {*s_i_* = 1, 2, …, *n*}. To maintain generality in our description, we consider a general observation error model (1), with

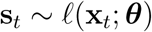

and with the latent movement path x_1:*T*_ defined by (11)-(12). We treat our discrete-time approximation (11)-(12) as the statistical model for the latent movement process, rather than the RSDE in (8)-(9). This requires a sufficiently fine temporal resolution *h* to maintain fidelity to the RSDE (8)-(9). Our goal is inference on the latent discrete-time representation of the animal’s movement path {x*_r_*, *r* = 1, 2, …, *T*} together with the movement parameters *θ* ≡ (*β*, *σ*^2^)′.

The main difficulty in such inference is the latent unknown movement path x_1:*T*_, because if we were able to condition on x_1:*T*_, inference on *θ* would be straightforward. If the latent movement path is unconstrained, then the model (1), (11)-(12) is a hidden Markov model (HMM), and inference can be made using recursive algorithms such as the Kalman filter (Cappé, 2005; Zucchini and MacDonald, 2009; Cressie and Wikle, 2011).

The projection in (12) is nonlinear, thus we need to make inference on the states and parameters in a nonlinear (constrained) state space model. Many methods for such inference have been proposed, including the ensemble Kalman filter (Katzfuss et al., 2016), MCMC (Cangelosi and Hooten, 2009), and particle filtering (Andrieu et al., 2010; Cappé et al., 2007; Del Moral et al., 2006; Kantas et al., 2009). In particle filtering, the filtering densities *f*(x*_t_*|s_1:*t*_; *θ*) are recursively approximated using particles that are propagated at each time point using the transition density (11)-(12), and then reweighted based on the observation likelihood *ℓ* (1). Particle filtering approaches to inference, like particle MCMC (Andrieu et al., 2010), are appealing for constrained processes because they do not require the evaluation of the transition densities (11)-(12), which are intractible due to the projection, but only require that they be simulated from.

### 3.1 Inference on RSDEs through Markov Chain Monte Carlo

To make inference on model parameters *θ* ≡ (*β*, *σ*)′ and the individual’s latent path x_1:*T*_, we constructed an MCMC algorithm to sample from the posterior distribution of x_1:*T*_, *θ*|s_1:*n*_. In doing so, we make explicit use of the simulation procedure in (11)-(12), which is a discretized, constrained movement model. It would be difficult to directly obtain the transition density function, because this would require marginalizing over the auxiliary 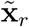

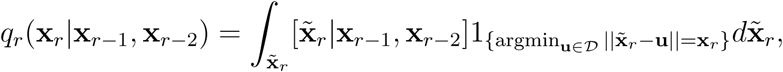

where 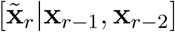 is given by (11).

However, because we have a tractable conditional density for 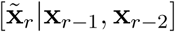, and x*_r_* is a deterministic function of 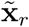 (x*_r_* is the projection of 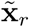 onto 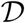), we constructed an MCMC algorithm that jointly updates 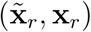, as follows. At the mth iteration of the MCMC algorithm, let the current state of the latent constrained process be 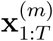, augmented by the unconstrained 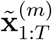. To update 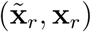 at one time point *r*, we propose a new location 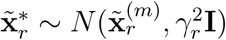 as a random walk centered on 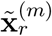, with proposal variance 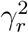. Projecting this proposed location onto 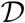 (if 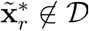) as in (12) gives 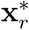, the proposed individual location at time *τ_r_*. The proposed pair 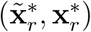 can then be accepted with probability

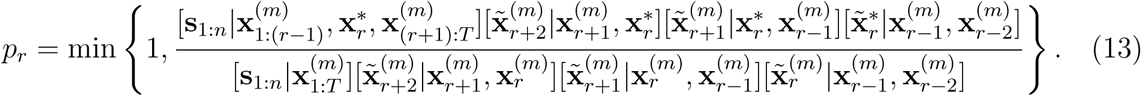

The likelihood of the data 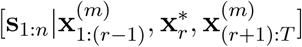 is the likelihood (1) of all observed telemetry locations, conditioned on the latent path and the proposed location 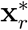 (e.g., Gaussian error for GPS data). Each of the transition densities (e.g., 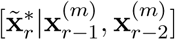) are multivariate normal densities given by (11).

This approach allows for Metropolis-Hastings updates for the latent locations x*_r_* one at a time. Block updates based on the simulation procedure could also be constructed. We found that updating each location at a time, using adaptive tuning (e.g., Craiu and Rosenthal, 2014) for each proposal variance 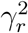 resulted in acceptable mixing, both in simulation (see Appendix B), and for the sea lion analysis in Section 4.

To complete the MCMC algorithm, we update the movement parameters *θ* ≡ (*β*, *σ*)′, conditioned on 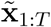. One could specify conjugate priors (e.g., a Gaussian prior for *β* and an inverse gamma prior for *σ*^2^), or one could use block Metropolis-Hastings updates to jointly update the movement parameters *β* and *σ* at each iteration of the MCMC algorithm. We favor this approach because it allows for more flexible prior specification. Similar update schemes could be used for parameters in the observation error model (1).

In Appendix B, we show a simulation example where we simulate movement constrained to lie within a polygon 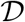, and make inference on model parameters. Code to replicate this simulation study is available upon request.

## 4 Modeling Constrained Sea Lion Movement

Having specified an RSDE-based approach for simulating animal trajectories constrained to lie within a domain 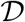, and for making inference on model parameters, we now apply this approach to the sea lion telemetry data.

### 4.1 Telemetry Data

As described in the Introduction, we consider the telemetry observations obtained from a sea lion off the coast of Alaska from December 6, 2010 to January 5, 2011. In this 30-day period of observation, *n* = 211 telemetry observations were obtained using the ARGOS system (ARGOS, 2015). The ARGOS system is unable to obtain a location fix when the sea lion is under water, thus the telemetry observations s_1:*n*_ were obtained at *n* irregular times *τ_i_*, *i* ∊{1, 2, …, *n*}. Each ARGOS telemetry location is also accompanied by a code *c_i_* ∊{3, 2, 1, 0, *A*, *B*} specifying the precision of the location fix at each time point, where *c_i_* = 3 corresponds to observations with the highest precision and *c_i_* = *B* corresponds to observations with the lowest precision. Multiple studies have shown that ARGOS error has a distinctive X-shaped pattern (Costa et al., 2010; Brost et al., 2015), and that each error class exhibits increasing error variance.

### 4.2 Model and Inference

To model the X-shaped error distribution, we follow Brost et al. (2015) and Buderman et al. (2016) and model observation error using a mixture of two multivariate t-distributed random variables, centered at the individual’s true location 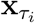

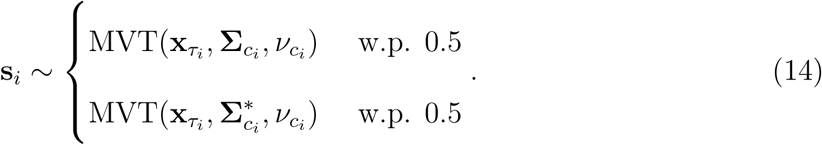

In (14), 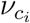 is the degrees of freedom parameter for ARGOS error class *c_i_*, and 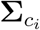 and 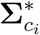 capture the X-shaped ARGOS error pattern

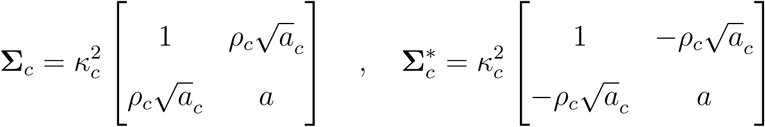

and *θ_c_* ≡ (*κ_c_*, *ρ_c_*, *a_c_*, *v_c_*)′ are ARGOS class-specific error parameters. See Brost et al. (2015) for additional details, and Costa et al. (2010) for an empirical analysis of ARGOS error patterns. As the distribution of ARGOS error has been studied extensively, we consider the ARGOS error parameters *θ_c_* to be fixed and known. For this study, we set these parameters equal to the posterior means reported in Appendix D of Brost et al. (2015).

To model sea lion movement, which is constrained to be in water 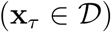, we consider a continuous-time model defined as a linear interpolation of the numerical approximation (11)-(12) of the RSDE (8)-(9). For a given temporal step size *h*, taken to be *h* = 5 minutes for this analysis, we consider an approximation to the RSDE at times *t_r_* ≡ *τ*_1_ + *rh*, *r* ∊{0, 1, …, *T*, *T* ≡ 30 × 24 × 12 = 8640}. At any observation time *τ_i_*, the individual’s position is given by a linear interpolation of the discrete approximation to the RSDE at the two nearest time points *t_r_*_(*i*)_, *t_r_*_(*i*)+1_, where *τ_i_* ∊ (*t_r_*_(*i*)_, *t_r_*_(*i*)+1_) and

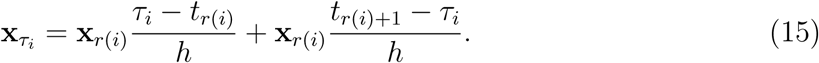

The RSDE model for movement is approximated at discrete times *t_r_* ≡ *τ*_1_ + *rh*, *r* ∊ {0, 1, …, *T*} according to (11)-(12). An alternative to this linear interpolation is to augment the approximation times (*t_r_* ≡ *τ*_1_ + *rh*, *r* ∊ {0, 1, …, T}) with the observation times (*τ_i_*, *i* = 1, …, *n*). This results in a non-uniform step size between time points at which the RSDE is approximated. The computational complexity of simulating the RSDE is linear in the number of time points, so the addition of the *n* additional time points is computationally feasible in many situations. The error in the numerical approximation (7) to the SDE scales with the largest time *h* between approximation times (Kloeden and Platen, 1992), so it is not clear that adding these *n* additional time points would result in an increase in numerical efficiency. We thus retain our regular temporal resolution, with a step size of *h*.

For this study of sea lion movement, we characterize space use over time. While it would be possible in some situations to model sea lion movement as being attracted to a haul-out or other central point (Hanks et al., 2011; Brost et al., 2015), we do not consider this here because our goal is only to characterize space use. Thus, we set *μ*(x*_τ_*, *τ*) = 0. We also assume a constant variance in the velocity process over time and space, with *σ*^2^ ≡ *c*^2^(x*_τ_*, *τ*). The resulting model for the discretized movement process constrained to be within water is

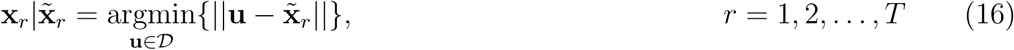

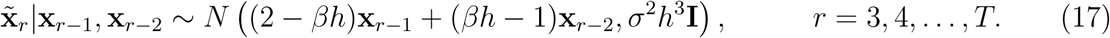

This model is illustrated conceptually in Figure 2.

**Figure 2:**
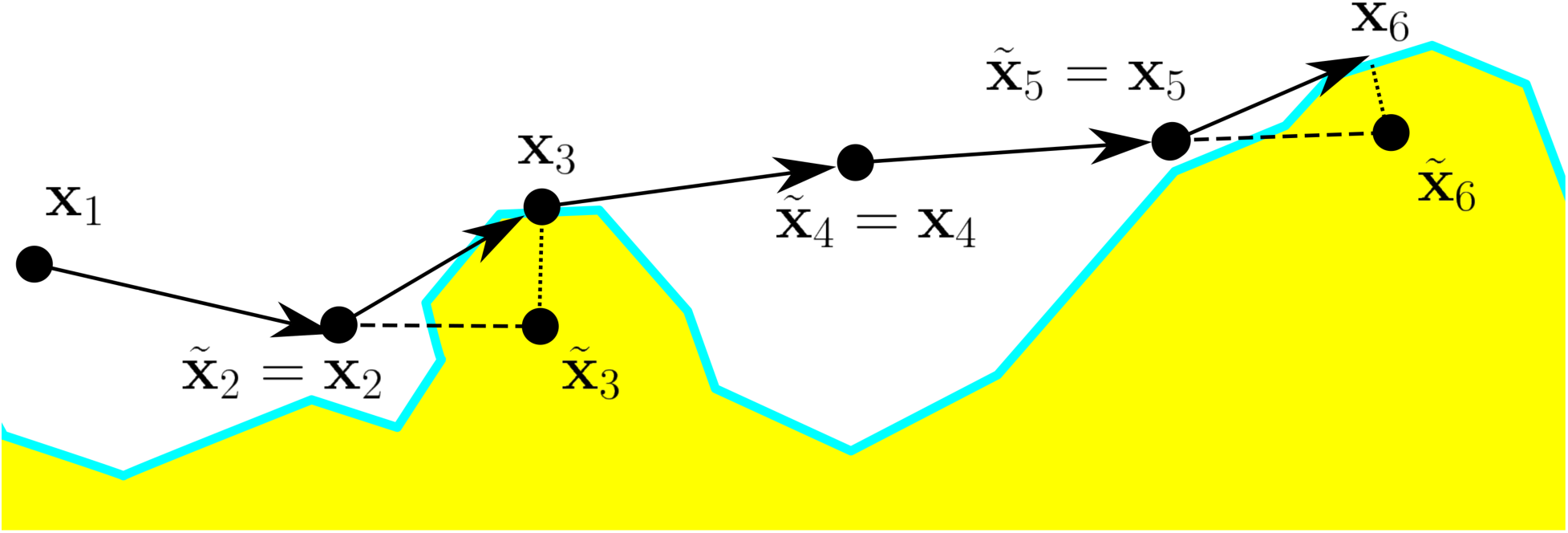
Modeling movement using projected processes. A time-discretized solution to the reflected SDE is obtained by first forward simulating from the transition density to obtain 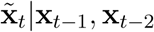, and then projecting 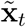 onto 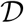 to obtain x*_t_*.

We complete the hierarchical state space model by specifying prior distributions for all parameters. For the initial two time points, we specify independent uniform priors

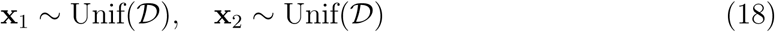

and we specify independent half-normal priors for the autocorrelation parameter *β* and the Brownian motion standard deviation *σ*

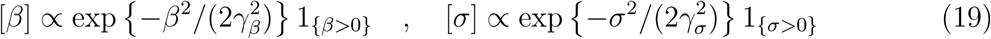

with *γ_σ_* = *γ_β_* = 100 as hyperparameters.

Our goal is inference on all parameters in the hierarchical Bayesian model for animal movement in (14)-(19). We constructed an MCMC algorithm to draw samples from the posterior distribution of model parameters, conditioned on the observed telemetry data, using methods described in Section 3.1. We used variable at a time Metropolis-Hastings updates (13) for the latent locations 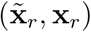, and used block Metropolis-Hastings updates to jointly update the movement parameters *β* and *σ* at each iteration of the MCMC algorithm. Random walk proposal distributions were specified for all parameters, and the variance of each proposal distribution was tuned adaptively using the log-adaptive procedure of Shaby and Wells (2010).

To initialize the MCMC algorithm for the sea lion analysis, we first chose starting values for movement parameters (*β*_0_ = .001, *σ*_0_ = 1km) and used a particle filtering algorithm (Cappé et al., 2007; Kantas et al., 2009) to provide a starting movement path x_1:*T*_ constrained to be in water. We initialized 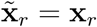 for each time point, and then ran the MCMC algorithm for 200,000 iterations. Convergence was assessed visually, with chains for *β, σ*, and each x*_r_* showing good mixing. The entire procedure required 14 hours on a single core of a 2.7GHz Intel Xeon processor. Code to replicate this analysis is available upon request.

### 4.3 Results

The posterior mean for log(*σ*), which controls the variance of the Brownian motion process on velocity, was 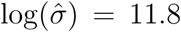, with an equal-tailed 95% credible interval of (11.2,12.4). The posterior mean for log(*β*), which controls autocorrelation beyond that implied by IBM, was 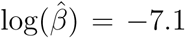, with an equal-tailed 95% credible interval of (–7.6, –6.2). The small estimated value for *β* implies that this term may not be needed in the model, and that IBM could be an appropriate model for this sea lion’s movement.

Figure 3a shows 10 realizations of paths x_1:*T*_ from the posterior distribution, and Figure 3b shows one path realization together with the observed ARGOS telemetry locations s_1:*T*_, with lines drawn from the telemetry locations to the realization of the individual’s location at the time of observation. Figure 4(a) shows 10 realizations from the posterior path distribution of the animal on December 11, 2010, as it navigated a narrow passage. Figures 4(b)-(j) show the posterior distribution of the sea lion’s location x*_t_* at 15 minute intervals. Our temporal discretization had a step-size of *h* = 5 minutes. Thus, there are two time points in our latent representation of the movement process between each shown time point in 4(b)-(j). Using a coarser time discretization would speed up computation at the expense of realism, as the linear interpolation (15) would result in paths that cross larger portions of land.

**Figure 3:**
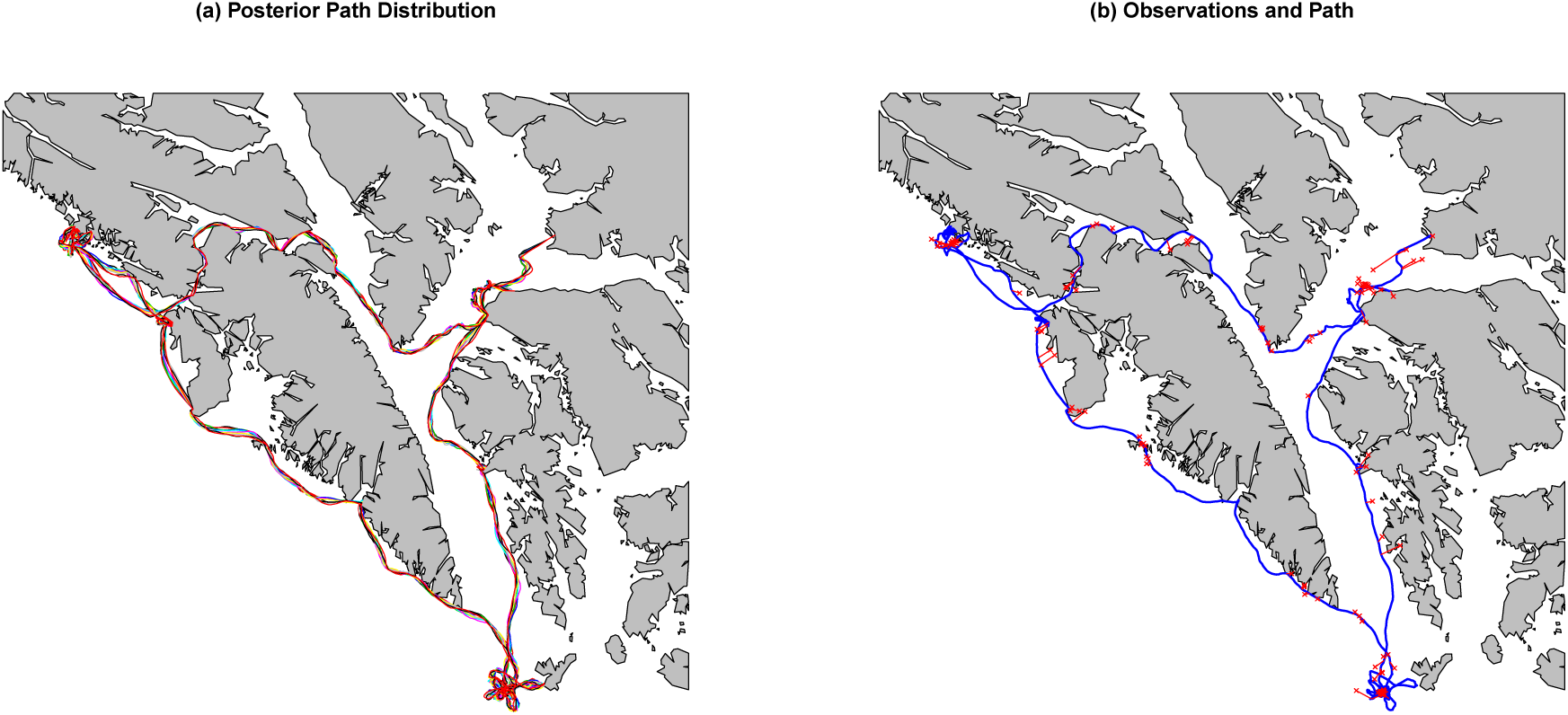
Ten sample paths from the posterior distribution of sea lion paths are shown in (a), with each path plotted in a different color. These paths show a propensity of the sea lion to stay close to coastlines. A single paths is shown in blue in (b), with the telemetry observations shown as red points, with lines connecting the telemetry observation to the estimated location 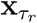. The imputed path between observed telemetry locations skirts islands and other barriers, as the movement model constrains the sea lion to be in the water or on the shoreline at all times.

**Figure 4:**
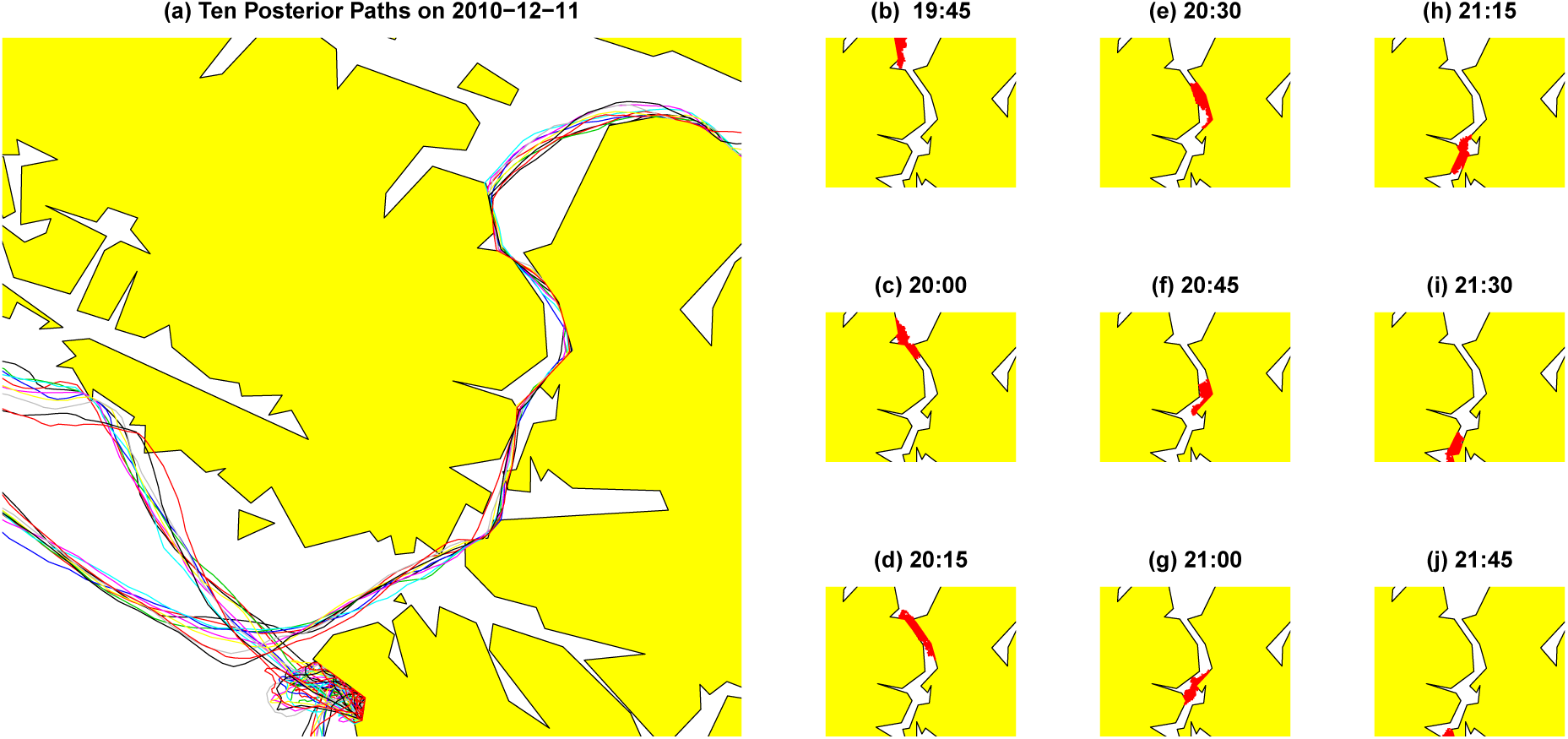
(a) Ten paths from the posterior distribution of sea lion locations as it navigates a narrow passage. (b)-(j) 5000 samples from the posterior distribution of sea lion locations at fifteen minute intervals. The posterior distribution shows that the individual is constrained to be within water (shown in white), or on the shoreline.

The spatial constraint is clear in both Figure 3 and Figure 4, and shows that our RSDE approach was successful in modeling realistic animal movement that is spatially constrained to occur within water (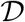). The posterior distribution of x_1:*T*_|s_1:*n*_ (Figure 3a) estimates sea lion space use over the 30 days of observation, and indicates that the individual spends a majority of its time near land, as expected for this species of pinniped.

## 5 Discussion

We developed an approach for modeling spatially-constrained animal movement based on a numerical approximation to a stochastic differential equation. The base SDE is general enough to capture a range of realistic animal movement, and the two-step procedure in Section 2.2.1 leads to a computationally tractable transition density. Our approach to constraining movement is based on the reflected SDE literature, and consists of projecting numerical approximations to the solution of the SDE onto the domain 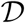.

Our approach for inference is computationally challenging, and future work will consider approaches that make inference more computationally efficient. The main computational burden for each iteration of the MCMC algorithm is projecting each latent 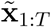 onto 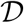. We coded Algorithm 1 using C++, but there is still significant room for improving computational efficiency. Algorithm 1 assumes that 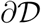 is given as a polygon or set of polygons, and checks each side of each polygon. When 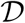 is large relative to the distance an animal can move between telemetry observations, Algorithm 1 could potentially be made more efficient by only considering a subset of polygon edges for each time point. An additional computational difficulty comes from the lack of conjugacy for the long latent time-series x_1:*T*._ Our approach is to use adaptively-tuned Metropolis-Hastings steps for each time point. One possible future approach is to construct a joint proposal for all 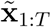 through a forward filtering, backward sampling algorithm (e.g., Cressie and Wikle, 2011) ignoring constraints. This would provide an approach for block updates of 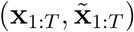 which may improve mixing of the MCMC algorithm.

#### Algorithm 1: Projected Solution to Reflected Stochastic Differential Equations

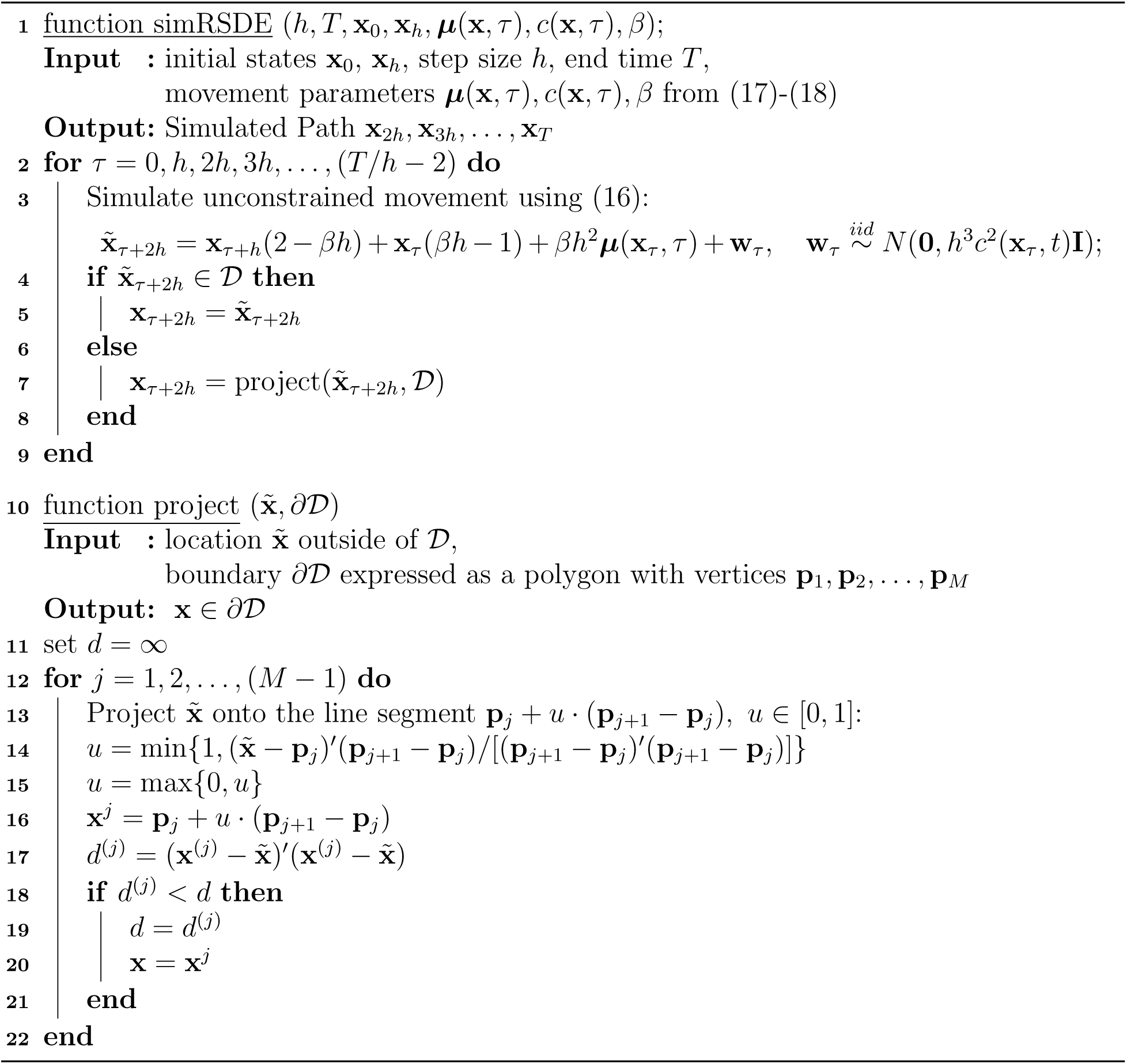

We note that there are other possible approaches to simulating RSDEs. One alternative, less common, approach for approximating the constrained SDE in (8)-(9) is to change the distribution of the increments (7) of the unconstrained numerical approximation to be distributed as truncated normal distributions with spatial constraint 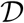 instead of the unconstrained Gaussian increments in (7). Cangelosi and Hooten (2009) used this approach and included a correction term in the mean of the truncated bivariate normal distribution to better capture the dynamics implied by the unconstrained SDE. Brillinger (2003) considered additional approaches to simulation, including specifying a mean function *μ*(x) that repels trajectories near the boundary 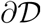. Russell et al. (2017) use a similar approach to constrain ant movement to lie within a nest.

Another approach to modeling movement constrained to lie within 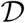 is to consider discrete space (gridded) approximations to the movement process (Hooten et al., 2010; Hanks et al., 2015; Avgar et al., 2016; Brost et al., 2015). A discrete support allows the spatial constraint on movement to be easily captured, but discrete space approaches can be computationally challenging to implement when the evaluation of the transition density requires the computation of all pairwise transitions from any grid cell to any other grid cell (e.g., Brost et al., 2015).

We assumed that animal locations are observed with error, which is the case for most telemetry data. If it can safely be assumed that observation locations have negligible error, then all observations will be within 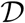. In this setting, the projection-based approach to inference developed in Section 3 could still be applied. This is true for SDE models that only model change in location, like potential function models (4), as well as SDE models that model change in velocity, like (2)-(3). An appealing alternative to the projection-based approach is to consider a truncated normal (TN) transition density with suitable location parameter *μ_t_*, covariance ∑*_t_*, and support 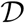

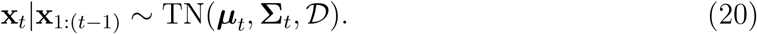

For example, the location and covariance parameters could be those defined by the approximation (11) to the SDE (8)-(9). The density function of the truncated normal distribution in (20) is

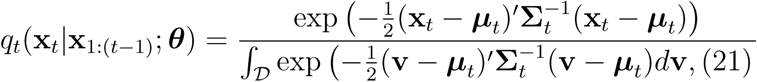

and fast approaches exist (Abramowitz and Stegun, 2012; Genz and Bretz, 2009) with accessible software (Meyer et al., 2016) for computing the normalizing constant when 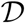 can be approximated as a polygon in 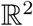.

We have focused on SDE models for the time-derivative of velocity. In some situations it is reasonable to model movement based solely on an SDE model for the time derivative of position (Brillinger et al., 2002; Preisler et al., 2013). The RSDE approach developed here could also be applied in this case, though there would be no need to do the two-step numerical approximation (7). Instead, an Euler approximation, or other numerical approximation to the SDE could be used (Kloeden and Platen, 1992).

Modeling movement without accounting for spatial constraints can lead to bias in movement parameter estimates and resulting inference. Our work, and the work of others who have also considered constrained movement (Brillinger, 2003; Cangelosi and Hooten, 2009; Brost et al., 2015) provide approaches that formally account for constraints and lead to more realistic animal movement and space use.

## Acknowledgments

Funding for this research was provided by NSF (DEB EEID 1414296), NIH (GM116927-01), NOAA (RWO 103), CPW (TO 1304), and NSF (DMS 1614392). Any use of trade, firm, or product names is for descriptive purposes only and does not imply endorsement by the U.S. Government. We thank Brett McClintock, Jay VerHoef, two anonymous reviewers, and an anonymous AE for their helpful suggestions on an earlier draft of this manuscript.

## APPENDIX A. Algorithm for Simulating RSDEs

We here describe an algorithm for simulating from RSDEs constrained to lie within a domain 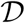.

## APPENDIX B. Simulation Example

In this Appendix, we consider a simple simulation example that illustrates the possible bias incurred by not accounting for constraints in movement. We consider a simple version of the reflected SDE in (8)-(9) in which *β* = 0 and *c*(x, *τ*) = *σ*

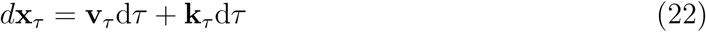

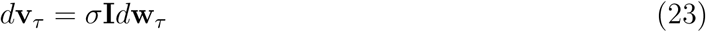

and where k*_τ_* is as given in (10). We refer to this constrained process as reflected integrated Brownian motion (RIBM) becuase the unconstrained version of this process (when k_*τ*_ = 0) is two-dimensional integrated Brownian motion (IBM). Figure B.1a shows a path simulated from RIBM for a given polygonal constraint 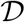. When the path is far from the boundary 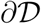, the path behaves identically to IBM. When the path is near the border 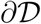, it often ends up identically on the boundary for short periods of time, because the minimal k process keeps x*_t_* from leaving 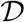. We also simulated noisy telemetry observations (Figure B.1b) under Gaussian observation error at 300 regularly spaced time points

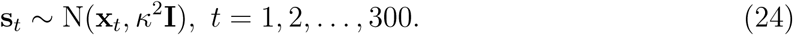

**Figure B.1:**
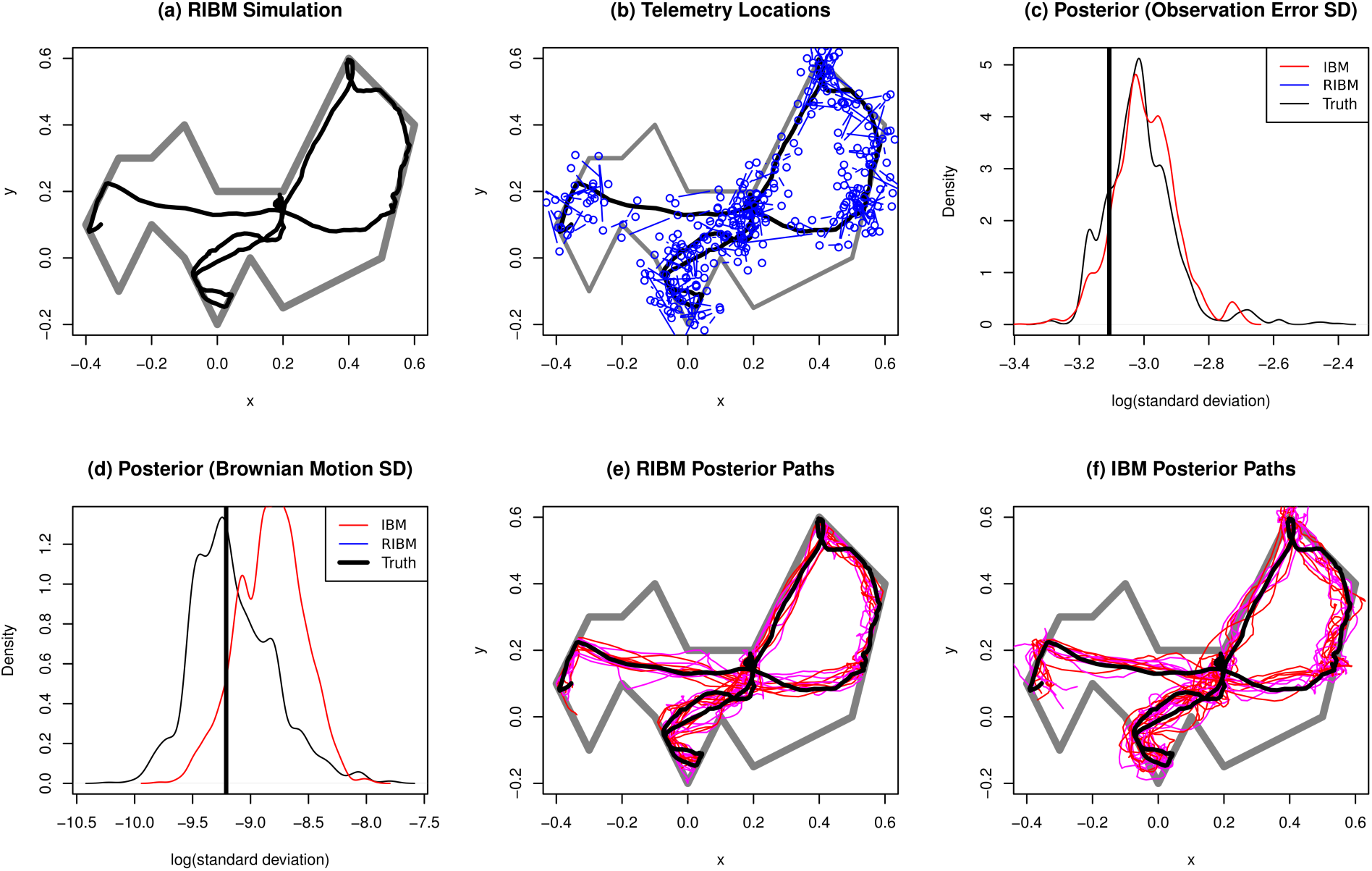
Simulation example (IBM). An individual’s path, constrained to occur within the polygon, was simulated using RIBM (a), and iid Gausian error was added (b) to simulate telemetry observation error. The observation error variance (c) and latent Brownian motion error variance (d) were estimated using PMMH for both RIBM and IBM models with diffuse half-normal priors. Results show some bias in the Brownian motion error variance. Samples from the posterior distribution of animal paths x_1:*T*_|s_1:*T*_ from the RIBM model (e) are constrained to lie within 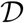, while those from the IBM model (f) are not.

We considered estimation of model parameters *θ* = (*σ*, *κ*)′ by specifying diffuse half-normal priors (with variance=100) for *σ* and *κ*, and sampling from the posterior distribution [*θ*|s1:300] using PMMH algorithms (Andrieu et al., 2010). Code to replicate this simulation study is available upon request. We considered estimation from the true model, RIBM, by constructing a particle filter using the projected simulation approach in Algorithm 1. We estimated model parameters using an unconstrained IBM model for x*_τ_* by constructing a particle filter without any projection or constraint. We note that it is possible to estimate *θ* under IBM by marginalizing over x_1:300_ using the convolution approach of Hooten and Johnson (2017), but we instead used PMMH to allow a more direct comparison between the estimates of *θ* under constrained (RIBM) and unconstrained (IBM) models. Each PMMH sampler was run for 10,000 iterations, with convergence of Markov chains assessed visually.

In general, the PMMH algorithm is less computationally-efficient for our system than the MCMC algorithm that we develop in the Section 3.1. The PMMH algorithm essentially attempts a block update of the entire latent path x_1:300_ at each MCMC algorithm, while the approach in Section 3.1 considers updating each x_*t*_ one at a time. Code to compare both of these approaches is available upon request.

Figure B.1c shows the estimated posterior distributions for the observation error standard deviation *κ* under IBM and RIBM, and Figure B.1d shows the posterior distributions for the Brownian motion standard deviation *σ*. There is little difference between constrained (RIBM) and unconstrained (IBM) models in the posterior distribution of the observation error *κ* (Figure B.1c), but the IBM model overestimates the Brownian motion standard deviation *σ* (Figure B.1d). Figure A.1e shows 10 sample paths from the posterior distribution of x_1:300_ under RIBM, and Figure 1f shows 10 sample paths from the posterior distribution under IBM. From this simulation example, it is clear that parameter estimates obtained without accounting for constraints in movement can show bias under model misspecification, though even under misspecification the 95% equal-tailed credible intervals of all model parameters under IBM include the true values simulated under RIBM. This may indicate that estimates obtained by fitting unconstrained movement models may be useful, even when we know the underlying movement process is constrained.

